# Network-based framework for studying etiology and phenotype diversity in primary ciliopathies

**DOI:** 10.1101/2025.01.08.631887

**Authors:** E.M. Aarts, D.S. Laman Trip, R. Neatu, C.G. Martin, B. Riley, A. Kraus, A. Green, M.H. Al-Hamed, R.E. Armstrong, J.A. Sayer, R. Bachmann-Gagescu, P Beltrao

## Abstract

Recent advances in sequencing technologies have increasingly enabled the identification of genetic causes for human monogenic diseases. However, systematic understanding remains limited due to the rarity, genetic heterogeneity, and complex genotype-phenotype relationships of these diseases. Primary ciliopathies are a diverse group of rare disorders caused by variants in genes associated with the cilium, a cellular organelle involved in signaling during development and cell homeostasis. These genetic variants result in a wide spectrum of clinical phenotypes involving the brain, eye, kidney and skeleton. It remains unclear to what extent this phenotypic diversity can be attributed to the disease-causing genes and their specific roles in ciliary function. Here, we systematically compared human primary ciliopathies with each other and with mouse phenotypes by propagating known disease genes through a network of protein interactions. Network propagation improved the clustering of primary ciliopathies with shared clinical phenotypes and facilitated the identification of mouse phenotypes closely related to primary ciliopathies due to shared groups of proteins in the interaction network. By leveraging this phenotype-specific approach, we prioritized candidate genes for ciliopathies and identified likely pathogenic variants in *CEP43*, a novel candidate gene for human primary ciliopathies in three previously unsolved cases. This study demonstrates that network propagation enhances the genetic and phenotypic understanding of primary ciliopathies, aiding in the prioritization of candidate genes and identification of relevant mouse models for these rare disorders, and providing a framework for unraveling shared underlying mechanisms for other rare genetic diseases.

## Introduction

In recent years, decreasing sequencing costs have significantly advanced our understanding of human genetic disorders. However, the precise etiology and molecular mechanisms of many inherited diseases remain elusive. For monogenic diseases, the genetic cause is often identifiable, yet a systematic understanding of disease etiology and its associated phenotypes is frequently lacking. The difficulty lies in the rarity and genetic heterogeneity of these diseases together with difficulties linking the genotype to complex phenotypes. To uncover disease etiology and molecular mechanisms, many studies have employed network-based approaches [1–5]. While these approaches have predominantly targeted common disorders, recent applications have extended to rare disorders too [6–8].

Network-based approaches have been utilized in numerous ways to study pleiotropy [9], drug targets [3], disease genes [2, 10], and molecular mechanisms [6]. A widely adopted method within these approaches is network propagation, which is based on the guilt-by-association principle [11]. In network propagation, disease genes are mapped onto a network of molecular interactions, and information is propagated over the network to enhance our understanding of diseases. With this approach, it is assumed that groups of interacting proteins will tend to have similar cellular roles and result in similar disease phenotypes when mutated. The link between interacting proteins and disease can also help highlight the underlying physiological and pathophysiological molecular mechanisms.

One group of rare monogenic disorders exemplifying the need for systematic studies are primary ciliopathies. Ciliopathies are caused by variants in genes associated with a single organelle, the cilium. Known as the antennas of the cell, cilia are sensory organelles perceiving, transducing and regulating a large variety of signals, thereby controlling cellular behavior during development and cell homeostasis. Variants in ciliary genes can result in a wide spectrum of clinical phenotypes, ranging from single tissue to multi-systemic disorders with symptoms including polydactyly, skeletal dysplasia, brain malformations, retinal disease, or kidney cysts [12]. There is significant phenotypic and genetic overlap between different primary ciliopathy syndromes, with limited success for efforts to study genotype-to-phenotype associations [13–17]. Indeed, it remains unclear to what extent the causal genes and their ciliary functions can explain the phenotypic divergence.

In this study, we systematically investigate rare disease etiology and phenotype-related molecular mechanisms through network propagation. First, we compare human primary ciliopathies using network propagation scores to uncover underlying proteins associated with different clinical phenotypes. To expand the limited knowledge on rare diseases, we demonstrate the use of genetic information from non-human models, such as mice, for enhancing our understanding of ciliopathies. Mouse models have been used extensively to further understand genetic diseases and their molecular mechanisms [18, 19]. However, it can be challenging to determine which mouse phenotypes accurately reflect human diseases. We developed an approach to identify mouse phenotypes related to human diseases based on network propagation scores and enhance candidate gene prioritization by leveraging genetic insights from these mouse models. We combine this approach with ciliopathy-specific network propagation scores to predict ciliopathy-specific disease genes, resulting in the identification of pathogenic variants in the novel ciliopathy gene *CEP43* for three patients with ciliopathy phenotypes and previously unsolved genetic diagnoses. Overall, our approach enhances our understanding of ciliopathies by identification of relevant mouse models and ranking of candidate genes and provides a framework for studying other rare genetic disorders using network-based methods.

## Results

### Network-based similarity between ciliopathies with shared clinical phenotypes

To systematically study ciliopathies, we applied network propagation to cluster traits including human ciliopathies and mouse phenotypes, identify trait-related protein modules, and predict candidate genes (Fig. 1a). For network propagation, we used a previously developed molecular interaction network [9] that integrates both direct (i.e., physical) and indirect protein-protein interactions (PPIs) from Reactome [20], IntAct [21], SIGNOR [22], and STRING [23] (Fig. 1b). We first collected known disease genes for each human ciliopathy from the Open Targets Platform (OTAR) [24], which aggregates data from several sources such as ClinVar [25], Orphanet [26], and Genomics England PanelApp [27], and used them as seed genes for network propagation. An expert in the field curated gene-trait pairs for ciliopathies. In total, we collected 174 unique disease genes for 21 primary ciliopathies, with the number of genes ranging from 2 to 42 per ciliopathy (Fig. 1c, Supplementary Table 1). Most evidence was obtained from Genomics England PanelApp, Orphanet, ClinVar, and Gene2Phenotype [28] (Fig. 1d, Supplementary Table 2). Furthermore, we annotated ciliopathies with their main clinical phenotypes, including central nervous system (CNS) disease, polydactyly, renal disease, retinal dystrophy, skeletal dysplasia, and hearing loss (Supplementary Table 1). Both clinical phenotypes and disease genes overlap between ciliopathies, e.g., *CEP290* is involved in many ciliopathies [29]. Additionally, some ciliopathies are known to have subtypes with different clinical phenotypes, such as “Joubert syndrome” (JBTS), “JBTS with ocular defect”, “JBTS with Jeune asphyxiating thoracic dystrophy (JATD)”, and “JBTS with renal defect”, which we kept separate. In this study, we only linked each given disease gene to one of the subtypes of a disease when the disease had multiple subtypes. Additionally, we excluded non-ciliary genes for retinitis pigmentosa (RP), as this disorder can also be caused by many other cellular mechanisms.

**Figure 1:**
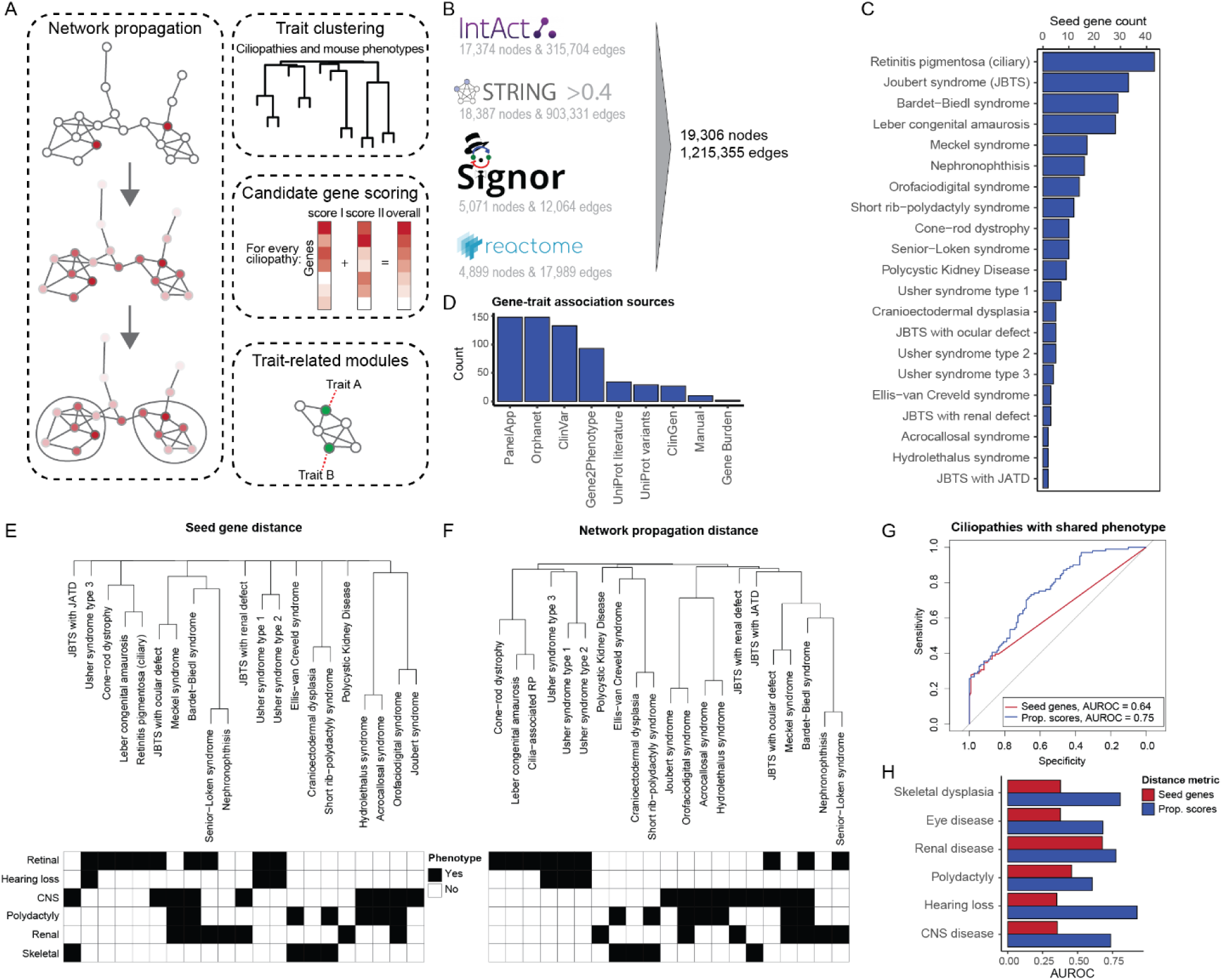
Network propagation improves phenotypic comparisons of ciliopathies. (A) Schematic of the approach: network propagation is used to compare traits (ciliopathies and mouse phenotypes), identify candidate genes, and detect underlying protein groups. (B) Number of nodes and edges from Intact, STRING, SIGNOR, and Reactome included in our undirected PPI network. (C) Number of disease genes for each of the 21 primary ciliopathies. (D) Number of gene-disease associations from each source of evidence. (E-F) Hierarchical clustering of ciliopathies based on overlap of known disease genes (Jaccard distance) (E) or network propagation scores (Euclidean distance) (F). Dendrograms are manually annotated with the main clinical organ phenotypes, including retinal disease, hearing loss, CNS disease, polydactyly, renal disease, and skeletal dysplasia. (G) ROC curves comparing recovery of ciliopathy pairs with at least one shared clinical phenotype based on known disease gene overlap (red) or on network propagation scores (blue). Grey line shows a random classifier. (H) AUC values for the predictions in (G), split by the clinical phenotypes. JATD, Jeune asphyxiating thoracic dystrophy.

To assess whether network propagation encodes phenotypic information on rare disorders, we performed network propagation for each ciliopathy, obtaining a score for every protein in the PPI network. This score reflects the distance of each protein to the studied disease and similar scores between distinct ciliopathies could suggest the involvement of similar proteins in the diseases. To analyze if ciliopathies with similar clinical phenotypes also have similar network propagation scores, we calculated the Euclidean distances between network propagation scores for all ciliopathies and created a hierarchical tree based on these distances (Fig. 1f). As a comparison, we calculated the overlap in known disease genes between the ciliopathies using the Jaccard distance to assess if we gain information from the network propagation scores (Fig. 1e, see methods). Clustering of ciliopathies using network propagation scores resulted in a hierarchical tree with more depth and branching, allowing for more comparisons between ciliopathies, as ciliopathies without shared disease genes could now be compared to other ciliopathies. To quantify this, we compared Jaccard and Euclidean distances between ciliopathies with at least one shared clinical phenotype and those without any shared phenotypes. We found that the distances from network propagation scores resulted in better recovery of ciliopathy pairs with a shared clinical phenotype, evaluated by the area under the ROC curve (AUROC) (AUROC = 0.75), compared to the recovery using the overlap of known disease genes (AUROC = 0.64, Fig. 1g). This suggests that genes associated with ciliopathies with shared phenotypes are connected within our PPI network.

Similarly, we found that the network propagation scores improved the recovery of ciliopathy pairs for each clinical phenotype individually (Fig. 1h). Network propagation mainly enhanced recovery of ciliopathy pairs with hearing loss, skeletal dysplasia, and CNS disease, and minimally enhanced recovery of ciliopathy pairs with renal involvement. The latter ciliopathies already had a high degree of shared genes, minimizing the value of the network propagation. These findings suggest that interconnected proteins may underlie the clinical phenotypes of ciliopathies, demonstrating a degree of phenotype specificity.

### Protein modules associated with primary ciliopathy phenotypes

To identify groups of highly connected proteins underlying the observed phenoype-specific clustering of ciliopathies, we clustered the PPI network into protein modules using Walktrap clustering [30] (Supplementary Table 3). We then linked individual ciliopathies to each protein module if the module contained at least one disease gene from the corresponding ciliopathy and was significantly enriched in high network propagation scores for that ciliopathy (adjusted p-value < 0.05, Wilcoxon Rank Sum test). Protein modules were annotated based on GO enrichment terms. We next focused on three protein module clusters assigned with ciliopathy-relevant GO terms related to cilia, visual process, and vesicle transport (Fig. 2a). We found that the visual protein modules were indeed mainly linked to eye-related ciliopathies, while protein modules in the “ciliary” cluster were linked to a broad range of ciliopathies. However, within the “ciliary” cluster, some specificity was observed. For example, the protein module “cellular response to potassium ion” contains several NIMMA related kinases (NEK) and Ankyrin Repeat And Sterile Alpha Motif Domain Containing (ANKS) proteins and is exclusively enriched for nephronophthisis. Another module designated with the GO term “BBSome, chaperonin-like” was only enriched for two ciliopathies with eye-related phenotypes (RP and BBS). Furthermore, the connectivity of disease genes in terms of protein interactions varied among ciliopathies. For instance, for some disorders such as cranioectodermal dysplasia (CED) or Usher type 1, the disease genes were found in a single module and were highly interconnected. In contrast, disease genes from Usher type 3 were distributed across multiple modules. This could suggest that Usher type 1 and CED are caused by more specific molecular mechanisms than Usher type 3.

**Figure 2:**
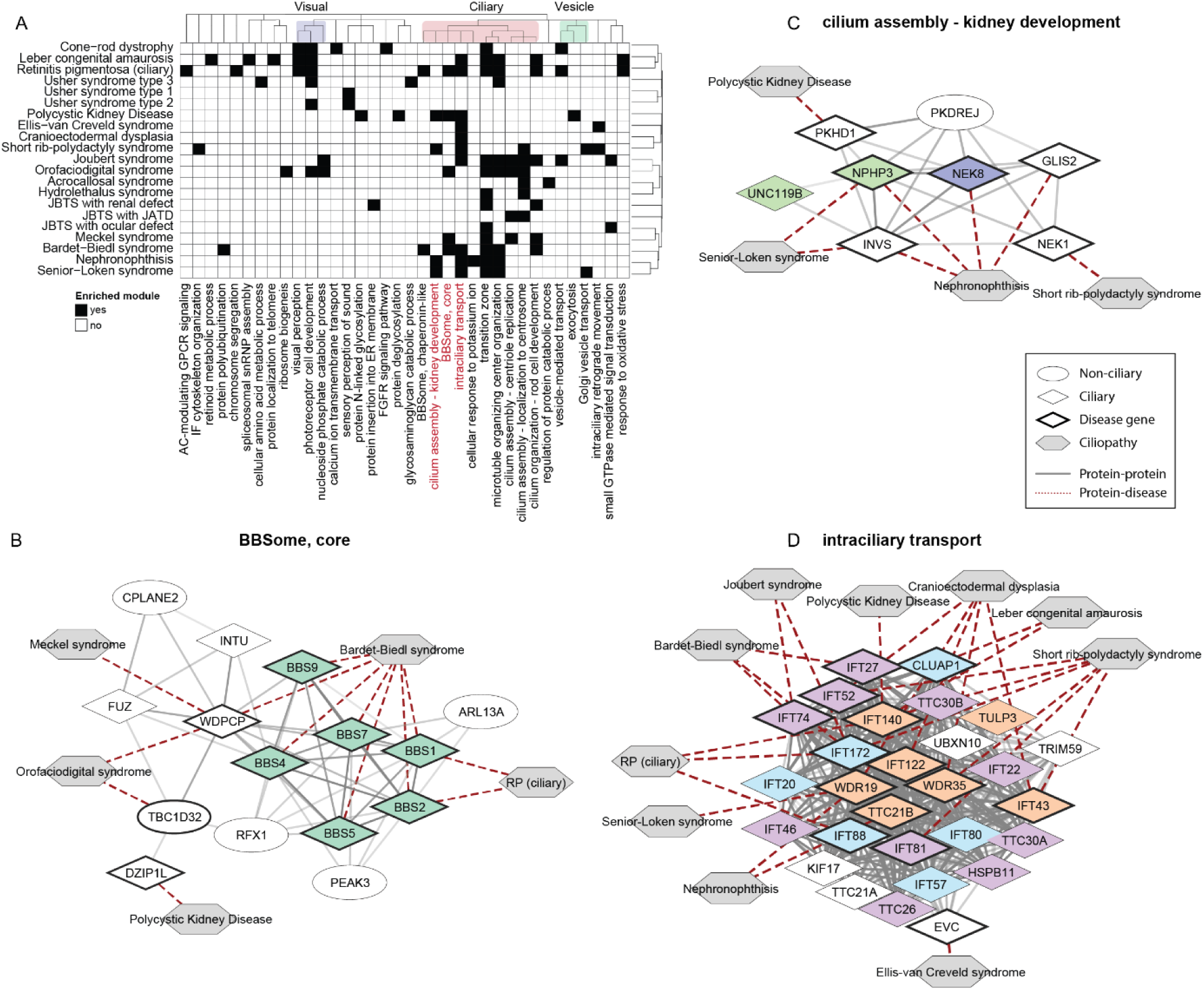
Identification of phenotype-specific protein modules in ciliopathies using network propagation scores. (A) Heatmap showing protein modules with significant enrichment of high network propagation scores for ciliopathies in black (adjusted p-value < 0.05, Wilcoxon Rank Sum test). Ciliopathies are clustered by network propagation scores and protein modules are clustered by re-clustering rounds of the network clustering. Protein modules describing similar functional terms are highlighted on top: “visual perception” (purple), “ciliary” (red), “vesicle” (green). Protein modules listed in red are detailed in the following panels. (B) Detailed view of the module “BBSome, core”. Genes in CiliaCarta [31] and SYSCILIA Gold Standard v2 (SCGSv2) [32] are represented by diamonds, other genes by ellipses, and ciliopathies by grey hexagons. Known ciliopathy genes are outlined in black (as based on initial determination of ciliopathy seed genes). Solid edges represent interactions from Intact or STRING, with transparency indicating the evidence score. Dotted edges indicate gene-disease associations. Nodes are colored based on selected complexes from Boldt et al. [33]: BBSome (green). (C) Detailed view of the module “cilium assembly – kidney development”. Node colors by Boldt et al.: ARL/NPHP/UNC119 (light green) and ANKS/NEK (blue). (D) Detailed view of the module “intraciliary transport”. Node colors by Boldt et al.: IFT-B1 (light blue), IFT-B2 (purple), and IFT-A (orange). AC, adenylate cyclase; GPCR, G protein-coupled receptor; IF, intermediate filament; FGFR, fibroblast growth factor receptor.

Looking into the protein modules, we found that some represented known protein complexes, such as the BBSome (Fig. 2b), associated with Bardet-Biedl syndrome (BBS). In addition to known BBSome complex proteins, the module also included proteins from the CPLANE complex, consisting of CPLANE2, INTU, FUZ, and WDPCP, and other proteins such as TBC1D32 and DZIP1L. Despite enrichment for BBS, we found no link to specific clinical phenotypes via single-organ disorders. In contrast, the ciliary protein module related to kidney function (Fig. 2c) was primarily linked to the renal ciliopathies Polycystic Kidney Disease (PKD), Senior-Loken syndrome (SLS) and NPHP. Lastly, intraciliary transport was associated with many ciliopathies (Fig. 2D). However, for ciliopathies with skeletal phenotypes, such as CED and SRP, most known disease genes can be found in this module (6/11 for SRP and 5/5 for CED), indicating that skeletal phenotypes are predominantly caused by changes in intraciliary transport.

### Comparison of network propagation between human ciliopathies and comprehensive mouse phenotypes

Having established that network propagation clusters together ciliopathies with similar phenotypes, we sought to apply the same methodology to identify mouse models that resemble human ciliopathies. Mouse models are chosen as they are extensively available for many genes and phenotypes. For example, the Mouse Genome Database includes 17,443 genes with targeted alleles and 74,265 genotypes with phenotype annotations [34]. By linking mouse models to ciliopathies through related genetics, this approach could provide mechanistic insight and an experimental platform for improving our understanding of rare human disorders.

We obtained all available 3,524 mouse phenotypes with their known associated genes from the Mouse Genome Database and propagated the associated genes for each phenotype through the protein interaction network independently (Supplementary Table 4). For each human ciliopathy, we then selected the twenty mouse phenotypes with the most similar network propagation scores as compared to the network propagation scores obtained for the human ciliopathies based on the Euclidean distances (see methods) (Supplementary Table 5). Finally, we clustered the selected mouse phenotypes and human ciliopathies using the network propagation scores and annotated the mouse phenotypes with their broader phenotype categories, termed ancestors, from the Mammalian Phenotype Ontology database [35] (Fig. 3a, Supplementary Table 6). We found that mouse phenotypes with similar ancestors generally cluster together. For example, we found that cluster 4 had a high fraction of vision/eye phenotypes while cluster 6 primarily contained skeletal phenotypes (Supplementary Fig. 1a). Moreover, we found that the mouse phenotypes often matched the clinical phenotypes of the human ciliopathies they clustered with. The vision/eye mouse phenotypes in cluster 4, for example, clustered together with eye-related human ciliopathies (Fig. 3b).

**Figure 3:**
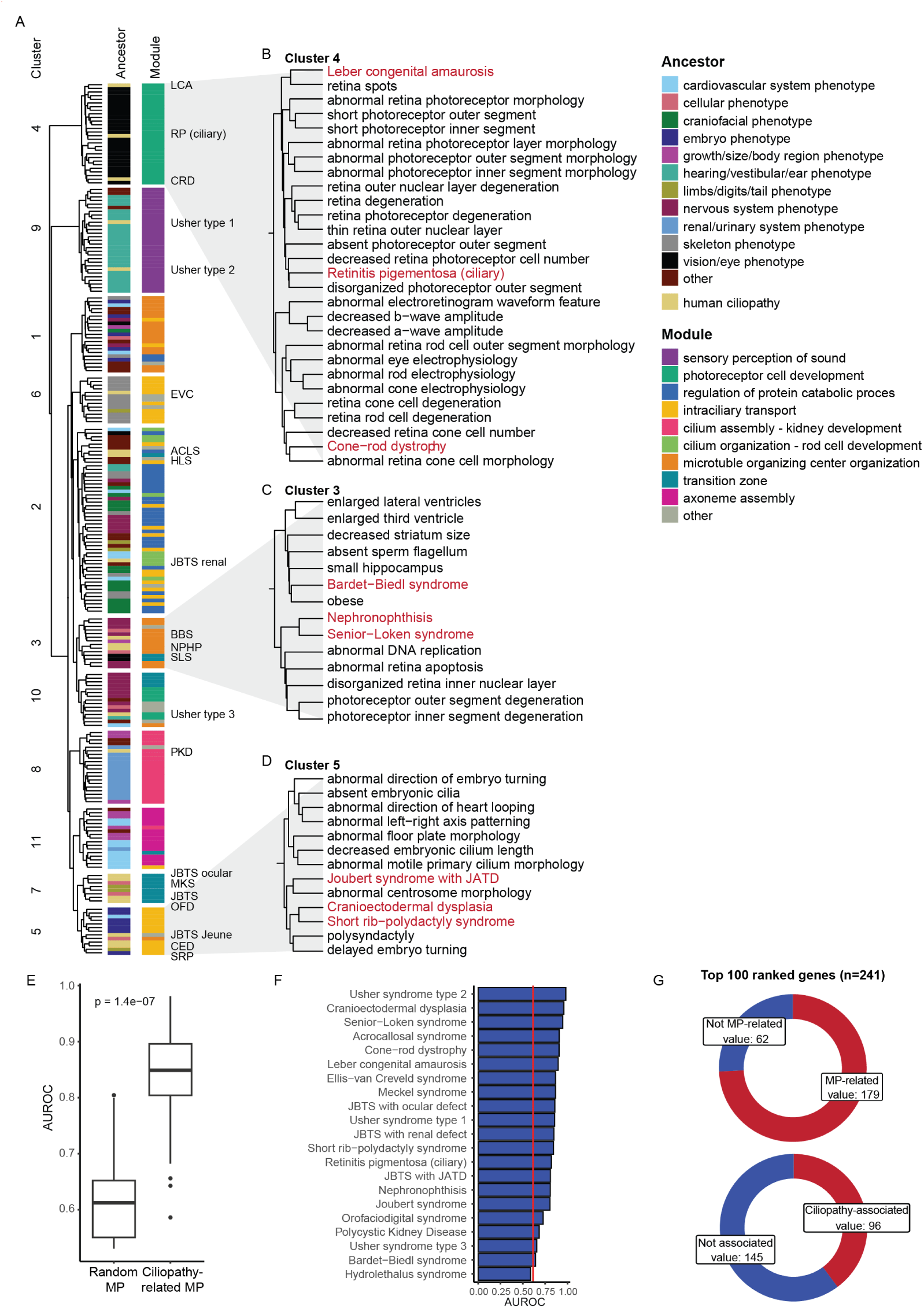
Identification of mouse phenotypes resembling individual ciliopathies through network propagation. (A) Hierarchical clustering of ciliopathies with related mouse phenotypes based on network propagation scores. Mouse phenotypes are annotated by their ancestor terms and all traits are annotated by their associated protein modules. For visualization, traits are color-based on the most represented ancestor term and protein module within each respective cluster. Rows containing human ciliopathies are labeled with their corresponding abbreviations and colored yellow in the ancestor annotation. (B-D) Zoom-in views of cluster 4 (B), cluster 3 (C), and cluster 5 (D). Human ciliopathies are in red and mouse phenotypes in black. (E) AUROC scores of known human ciliopathy gene predictions using network propagation scores from ciliopathy-related mouse phenotypes versus randomly selected mouse phenotypes. Ciliopathy disease genes were excluded from the seed genes of mouse phenotypes prior to running network propagation. Genes were ranked for each ciliopathy based on the combined network propagation scores of selected mouse phenotypes versus random mouse phenotypes. (F) AUROC scores from (E) visualized per ciliopathy. The red line marks the median AUROC scores when using random mouse phenotypes. G) Donut plots of the top 100 ranked genes for each ciliopathy by mouse phenotypes indicating whether genes were already associated to those mouse phenotypes (top) or to human ciliopathies (bottom). MP, mouse phenotype; LCA, Leber congenital amaurosis; RP, Retinitis Pigmentosa, CRD, cone-rod dystrophy; EVC, Ellis-van Creveld syndrome; ACLS, acrocallosal syndrome; HLS; Hydrolethalus syndrome; JBTS, Joubert syndrome; BBS Bardet-Biedl syndrome; NPHP, Nephronophthisis; SLS, Senior-Løken syndrome; PKD, Polycystic Kidney Disease; MKS, Meckel syndrome; OFD, orofaciodigital syndrome; CED, Cranio-Ectodermal Dysplasia; SRP, Short-Rib-Polydactyly syndrome.

In contrast, most multi-systemic ciliopathies clustered with mouse phenotypes with a high variety of ancestors. However, these traits were linked via the enrichment of their network propagation scores in the same protein modules (Supplementary Fig. 1b) (Supplementary Table 7). For instance, cluster 3 had a high fraction of both human and mouse traits with high network propagation scores in the module ‘microtubule organizing center organization’, containing proteins important for ciliogenesis, but included diverse mouse phenotypes, ranging from obesity to photoreceptor degeneration (Fig. 3c). Interestingly, these mouse phenotypes still partly resembled the clinical phenotypes of BBS (obesity, CNS disease, retinal disease), NPHP (retinal disease), and SLS (retinal disease). Similarly, cluster 5, enriched for the module ‘intraciliary transport’, connected skeletal ciliopathies together with a variety of mouse phenotypes, such as polysyndactyly and abnormal centrosome morphology (Fig. 3d). Overall, this demonstrates that comparing rare human disorders with mouse phenotypes using network propagation can identify mouse phenotypes resembling human disorders.

As we were able to identify ciliopathy-related mouse phenotypes using network propagation, we wondered if we could use the genetic information from these mouse models to propose novel human ciliopathy genes. To test this, we excluded known human ciliopathy genes from the mouse phenotype-associated genes and tried to retrieve the human genes using network propagation with the remaining mouse genes as seed genes (see methods). This worked significantly better when using ciliopathy-related mouse phenotypes compared to random mouse phenotypes (Fig. 3e, p-value = 1.4e-7; two sample t-test) and best when using the ten mouse phenotypes with the most similar propagation scores compared to including more phenotypes (Supplementary Fig. 1c). The effectiveness of gene recovery varied across human ciliopathies, with recovery of Usher type 2 genes performing the best and hydrolethalus syndrome (HLS) genes the worst (Fig. 3f). We found that the combination of mouse phenotypes for predicting human ciliopathy genes identified additional high-scoring genes that were not directly linked to the mouse phenotypes or human ciliopathies (Fig. 3g). For every ciliopathy, we took the top 100 ranked genes, which resulted in a total of 241 unique genes. Of these, 62/241 were not yet associated with the mouse phenotypes used in the analysis and 145/241 were not yet linked to a human ciliopathy. This suggests that integrating multiple mouse phenotypes allows for the discovery of additional potentially relevant disease genes compared to just taking the known genes associated with the separate mouse phenotypes.

### Disease-specific network propagation scores and disease-related mouse phenotypes accurately prioritize human ciliopathy genes

To further improve identification of novel candidate genes for ciliopathies, we combined mouse phenotype-based ranking of genes with disease-specific network propagation scores and gene expression patterns in human tissues. Here, we accounted for biases towards well-studied genes by permuting the network propagation scores (Supplementary Fig. 2). For the gene expression patterns, we obtained single-cell RNA sequencing data from the Human Protein Atlas (HPA) (proteinatlas.org) [36]. Since ciliary genes are typically expressed at low levels, we used relative gene expression to identify cell types with the highest and lowest expression of known ciliopathy genes (Supplementary Fig. 3a). Cell types, as defined by HPA, with high expression included photoreceptor cells, astrocytes, spermatocytes, and ciliated cells. By comparing the expression patterns of all genes to those of known ciliopathy genes (see methods), a logistic regression model successfully retrieved ciliopathy genes that were removed for testing (Supplementary Fig. 3b) (AUROC = 0.79).

To integrate the three scores for each ciliopathy (permuted network propagation, mouse phenotype ranking, and expression pattern), a logistic regression model was trained on four ciliopathies with the highest numbers of known genes (RP, JBTS, BBS, and LCA). Because of the limited number of known genes per ciliopathy, we trained the model parameters for the four ciliopathies together. However, the predictions are still ciliopathy specific as the feature scores are calculated separately for each ciliopathy.

To test the model, we selected six ciliopathies not used for training with at least ten known disease genes per ciliopathy. Combining the permuted network propagation scores with the mouse phenotype ranking significantly increased the precision-recall AUC (PR AUC) (Fig. 4a) and partial AUROC (0.95-1.0 specificity) (Fig. 4b), compared to the separate scores alone. We found limited differences for the AUROC, likely due to the small fraction of positive data points (Supplementary Fig. 4a). We observed performance differences between the different ciliopathies, but combining the two scores yielded the best result in most cases (Supplementary Fig. 4b). Interestingly, the gene expression score did not improve the model, potentially due to its generalization across all ciliopathies. For predicting candidate genes for each ciliopathy individually, our gene expression model was not specific enough. Therefore, we proceeded with the model only combining the permuted network propagation and mouse phenotype ranking features (Supplementary Table 8).

**Figure 4:**
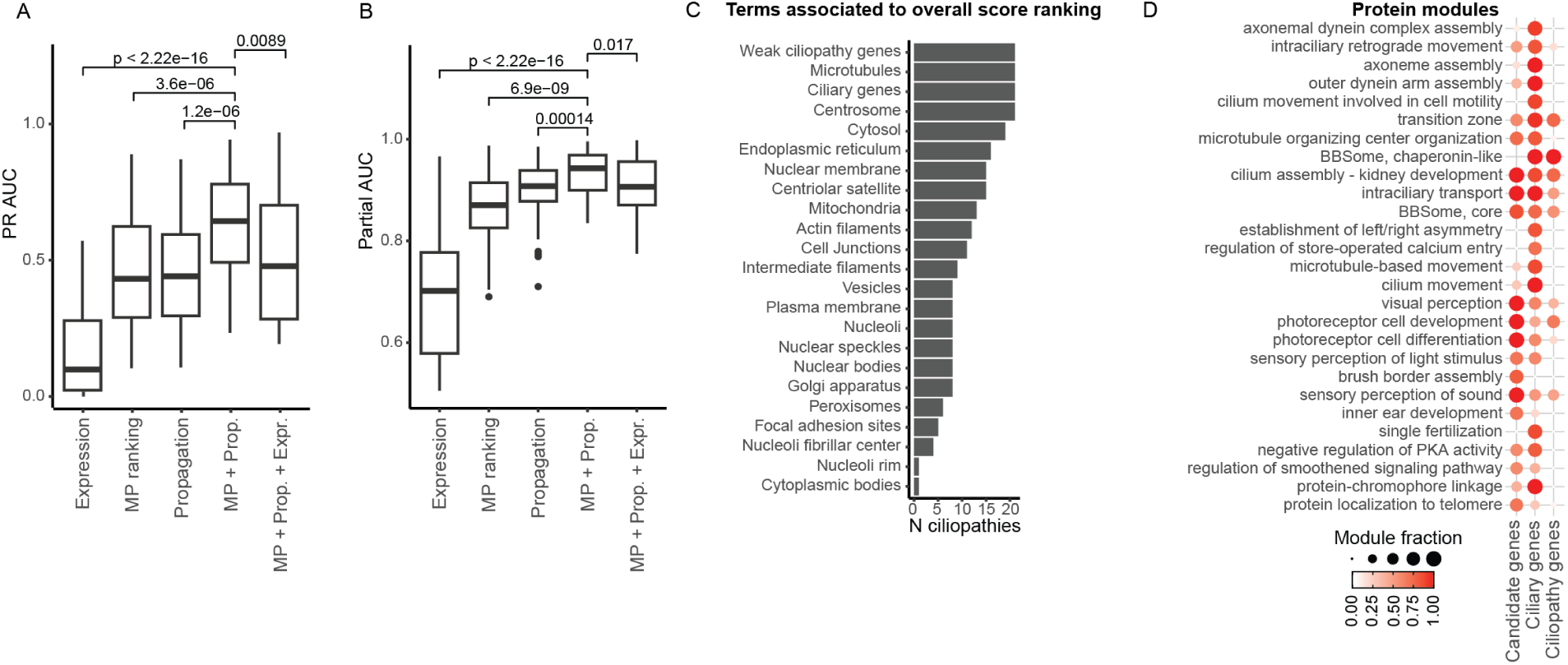
Model for candidate gene prediction specific to each ciliopathy. (A) PR AUC for logistic regression models combining mouse phenotype ranking (MP), permuted network propagation scores (Propagation), and gene expression scores (Expression) in various combinations. B) Partial AUC (1.00-0.95 specificity) for the same models as in (A). (C) Number of ciliopathies (N ciliopathies) with significant GSEA for defined gene sets. Weak ciliopathy genes = genes from databases with low confidence scores. Ciliary genes = genes in CiliaCarta and SCGSv2. Other gene sets are obtained from Human Protein Atlas (HPA) localization data. (D) Fraction of genes in protein modules annotated as candidate, ciliary gene, or known ciliopathy gene. Only protein modules containing a fraction of candidate genes, ciliary genes, or ciliopathy genes above 0.6 are shown. Prop., propagation; Expr., expression; PKA, protein kinase A.

To analyze which genes were highly ranked using our model, we performed gene set enrichment analyses (GSEA) for several gene sets: genes associated with ciliopathies in databases but with low confidence scores (termed “weak ciliopathy genes”), known ciliary genes (CiliaCarta and SCGSv2), and protein localization data from HPA. High-ranking genes for all ciliopathies were enriched for “weak ciliopathy genes”, known ciliary genes, and proteins localized to microtubules and centrosomes, suggesting that our model effectively predicted cilia-related genes (Fig. 4c). However, not all ciliary genes were highly ranked, indicating that our model makes some separation between those ciliary genes likely involved in ciliopathies and those ciliary genes that are not (Supplementary Fig. 5a). In addition, the model preserved phenotype specificity, as gene rankings for related ciliopathies were correlated (Supplementary Fig. 5b).

We also investigated whether our identified protein modules described above were enriched for known ciliopathy genes, candidate genes (top 100 per ciliopathy), or ciliary genes (Fig. 4d). Most modules containing candidate genes had a high proportion of known ciliopathy genes. However, we also identified some modules containing candidate genes that had low fractions of known ciliopathy genes, such as those related to localization to telomeres, protein kinase activity, and Hedgehog signaling. Additionally, we identified modules with a high fraction of ciliary genes but a low fraction of known or candidate ciliopathy genes, such as those related to calcium entry and left/right asymmetry. The most promising candidate genes are likely within modules with a high fraction of known ciliopathy genes, as these modules are already linked to ciliopathies. However, genes in other modules could reveal more novel biology.

### Network propagation identifies *CEP43* as novel candidate gene

Using our model, we generated a list of 285 unique candidate genes highly ranked for various ciliopathies (Supplementary Table 9). We next sought to identify ciliopathy patients with unsolved genetic diagnoses. First, we searched a cohort of 75 patients with BBS (38), JBTS (13), or multi-systemic ciliopathies (24) from the Genomics England 100,000 genomes database [37] for homozygous and compound heterozygous variants in the 285 candidate genes. In addition, we used variant effect predictors (VEPs), including CADD [38], ESM1b [39], AlphaMissense [40], and spliceAI [41] to estimate pathogenicity of the identified candidate gene variants.

We identified one patient with homozygous variants in *CEP43*, also known as *FGFR1OP* or *FOP*, which was ranked in the top 100 highest ranked candidate genes for multiple ciliopathies (Fig. 5a). A follow-up expanded search in the Genomics England database for individuals with bi-allelic *CEP43* variants identified one more individual with clinical diagnoses related to primary ciliopathies. In addition, a third individual was identified in a separate cohort. The three individuals had various diagnoses, including BBS, CRD, and skeletal dysplasia (Fig. 5b, Supplementary Table 10). CEP43 was found to interact with proteins already known to cause ciliopathies and ciliopathy-related mouse phenotypes (Fig. 5c). For example, CDK5RAP2 is known to be associated with a small hippocampus in mice and CEP135 with polydactyly, which are both BBS-related phenotypes.

**Figure 5:**
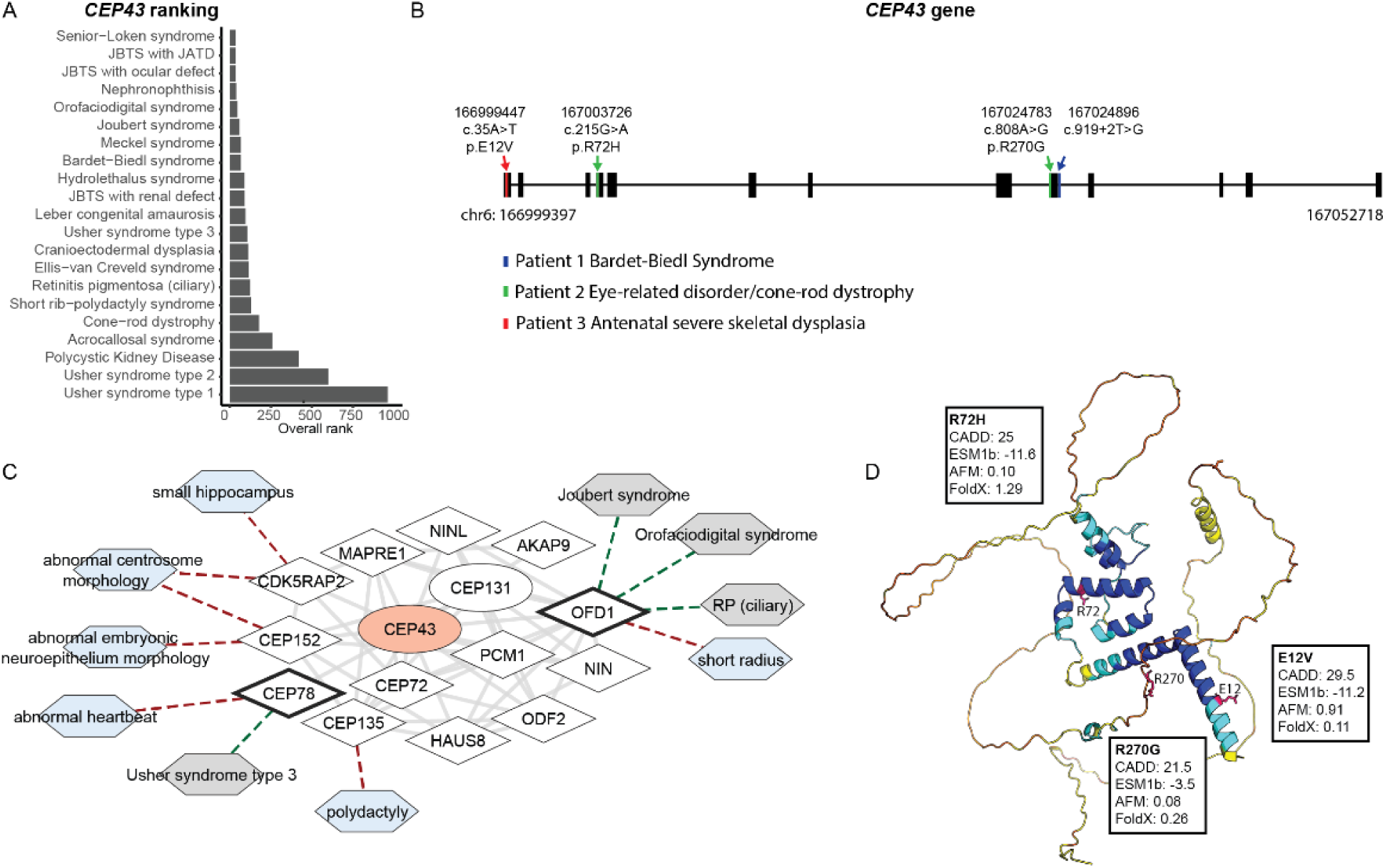
CEP43 as novel ciliopathy disease gene. (A) Overall rank of CEP43 for ciliopathies. (B) Schematic representation of the CEP43 gene, highlighting variants identified in three patients, annotated with their clinical diagnoses. Variants are homozygous in patients 1 (blue) and 3 (red). (C) Protein module including CEP43, filtered to show proteins directly connected to CEP43. Genes in CiliaCarta and SCGSv2 are represented as diamonds, other genes as ellipses, and traits as hexagons. Known ciliopathy genes have a black border and CEP43 is visualized in red. Mouse phenotypes are colored blue and human ciliopathies grey. Solid edges represent protein interactions from Intact or STRING, with transparency indicating evidence scoring. Dotted edges denote gene-trait connections (red to mouse phenotypes and green to human ciliopathies). (D) Predicted protein structure from the AlphaFold Database [46] of CEP43 showing coding variants from (B), annotated with their CADD, ESM1b, AlphaMissense, and FoldX scores.

The individual with a BBS-like phenotype carried a homozygous splice region variant c.919+2 (NC_000006.12:g.167024896T>G) with a CADD score of 33 and a high predicted impact (spliceAI delta score = 0.99) (Fig. 5b). The individual diagnosed with CRD was found to have two heterozygous variants in *CEP43* (NC_000006.12:g.167003726G>A and NC_000006.12:g.167024783A>G) (Fig. 5b). Both variants have high predicted CADD scores but only p.R72H (NC_000006.12:g.167003726G>A) is predicted to be pathogenic by ESM1b and destabilizing by FoldX (Fig. 5d), while p.R270G (NC_000006.12:167024783A>G) is predicted to be a splice acceptor variant by spliceAI (delta score = 0.34). Lastly, an individual with antenatal severe skeletal dysplasia (pregnancy terminated) was identified with a homozygous p.E12V variant (NC_000006.12:g.166999447A>T). This variant is predicted to be pathogenic by CADD, ESM1b, and AlphaMissense but not predicted to be destabilizing, suggesting another mechanism of pathogenicity.

In previous literature, *CEP43* has already been shown to be important for cilia assembly and disassembly and has a mouse model with a skeletal phenotype [42–45]. Here we now identify patients with a diverse range of primary ciliopathy phenotypes (retinal, skeletal and neurological) harboring pathogenic variants in this gene. Together, these observations demonstrate that our model combining network propagation scores with mouse phenotype rankings prioritizes candidate disease genes that are found to have variants in ciliopathy patients with unresolved genetic diagnoses.

## Discussion

Our study provides a framework for studying disease etiology and genotype-to-phenotype associations in rare diseases. As proof of principle, we show that network propagation scores contain phenotypic information for primary ciliopathies, as these scores improved retrieval of disease pairs with similar clinical phenotypes, compared to overlap in known disease genes. We also obtained insights into protein groups underlying these phenotypes upon clustering of the PPI network. Similarly, we show that the same approach can be used to identify mouse phenotypes related to rare diseases based on their genetics and that these similarities relate to the clinical phenotypes. Furthermore, genetic knowledge of these mouse phenotypes improved prediction of disease genes.

While ciliopathies are characterized by dysfunction of proteins linked to one organelle, there is a large variety in their clinical phenotypes. As ciliopathies are rare and genetically heterogeneous, it is challenging to assess how much phenotypic diversity can be explained purely by the disease-causing genes themselves. In our analysis, we have identified phenotype-specific protein modules which may facilitate the identification of ciliary pathomechanisms linked to specific clinical phenotypes. For primary ciliopathies, specificity was found in protein modules related to visual, hearing, renal, and skeletal phenotypes. These organ-specific protein modules also contained genes associated with multi-systemic ciliopathies. This could suggest that individuals with multi-systemic ciliopathies and mutations in genes within these modules are more likely to develop involvement of the corresponding organ. For instance, the module ‘cilium assembly – kidney development’ was enriched for ciliopathies with a renal phenotype, including NPHP, PKD, and SLS. Almost half of the mouse phenotypes enriched for this module were also related to the renal/urinary system. In addition, the skeletal ciliopathy SRP was associated with this module via Nek1. As a Nek1 mouse model has a renal phenotype [47] and some SRP subtypes also have renal involvement, this can suggest that patients with *NEK1*-associated SRP may be more likely to develop kidney disease than those with SRP caused by other genes. Such suggested associations must now be investigated in human ciliopathy cohorts, whereby the small number of affected patients may limit the analysis.

For genotype-to-phenotype associations, we were also particularly interested in JBTS as multiple clinical subtypes of this disease have been described. When including the mouse phenotypes, JBTS with the skeletal phenotype JATD clustered together with the skeletal ciliopathies CED and SRP, indicating similar underlying mechanisms. One possibility to explore could be an abnormal centrosome morphology, as mice with this cellular phenotype cluster together with the skeletal human ciliopathies. The subtypes of JBTS with renal defect and JBTS with ocular defect did not specifically cluster with clinically similar human ciliopathies or renal/ocular mouse phenotypes, which could indicate that the current known subtype-associated genes are not specific enough to the clinical phenotype. Still, proteins associated with pure JBTS but clustered with proteins causing a specific subtype might be interesting targets for further studies into JBTS subtypes. For example, the protein modules related to centrosome localization and centriole replication contain proteins associated with pure JBTS and JBTS with JATD, suggesting that genes in this module, known to cause pure JBTS – such as *CEP120*, *KIAA0753*, *KIF7*, and *PIBF1* – may also cause JBTS with JATD.

In addition to organ-specific protein modules, we also identified protein modules associated with a large variety of human ciliopathies and mouse phenotypes. For instance, the module termed ‘BBSome, core’ was associated with various human ciliopathies including BBS but could not be linked to a specific clinical phenotype. However, these protein modules can still give more insights into molecular mechanisms. We identified a strong connection between proteins from the CPLANE complex, consisting of INTU, FUZ, WDPCP, and CPLANE2, and the BBSome core proteins. *WDPCP* has already been identified as a BBS gene [48], but the other proteins have not been clearly linked to the BBSome. A recent study did show that FUZ might link GPR161 to the BBSome for removal from cilia [49], also suggesting a link between FUZ and the BBSome. In further studies, the links between the CPLANE complex, other proteins identified here in a BBSome core protein module, such as RFX1, TBC1D32, and ARL13A, and the BBSome should be investigated.

Furthermore, we showed that genetic knowledge can be transferred from other groups of traits to rare diseases. It has already been shown that GWAS traits share genes with ciliopathies (42) and that mouse models can help ciliopathy gene prediction [19]. Here, we demonstrate that network propagation scores can systematically rank mouse phenotypes based on their relevance to the corresponding rare disorder. Disease gene retrieval based on mouse phenotypes was not equally accurate for all human ciliopathies, suggesting that some high-ranked mouse phenotypes are related to the corresponding human ciliopathy primarily due to shared known genes, while other genes associated with the mouse phenotypes do not provide additional insights into human ciliopathies. Interestingly, the retrieval of human ciliopathy genes using randomly selected mouse phenotypes for prediction still outperformed random gene selection, suggesting a potential bias in either the available mouse models or the network. Combining information from mouse phenotypes with network propagation scores was shown to accurately predict ciliopathy-associated genes, resulting in the identification of likely pathogenic variants in *CEP43*, also known as *FGFR1OP* or *FOP* in three individuals with primary ciliopathy syndromes including a BBS-like phenotype, a severe skeletal dysplasia, and a retinal phenotype.

Even though our study is helpful in prioritizing ciliopathy candidate genes, the identification of novel ciliary genes in general might be limited. As our analysis is based on known protein interactions, the bias of current research will be reflected in our results. In addition, databases of disease genes are not always up to date compared to the literature. Therefore, some candidate genes from this study may have already been shown to cause ciliopathies in recent papers, such as *SCLT1* [51] and *FUZ* [52]. Lastly, our analysis is limited by the generalization of the interactome. Tissue- or cilium-specific interaction networks might improve our analysis and prevent false positive interactors in some tissues [53].

Overall, we performed a systematic analysis of rare disease etiology and phenotype-specific protein modules in primary ciliopathies. We identified mouse phenotypes that give a good representation of ciliopathies, which can be used for further studies of disease mechanisms. We also employed these models to prioritize ciliopathy-specific candidate genes, and we identified novel likely pathogenic variants in three ciliopathy patients. In this study, we focused on ciliopathies, but this approach can be used to study etiology and genotype-to-phenotype associations in a large set of rare disorders.

## Methods

### Network propagation for ciliopathies and mouse phenotypes

For the PPI network, we used the Open Targets Interactome Network (release 22-07-2022). This network integrates direct (i.e., physical) and indirect interactions from IntAct, SIGNOR, Reactome, and STRING (scores >= 0.4). Duplicated edges and self-loops were removed, resulting in a network containing 19,306 nodes and 1,215,355 edges.

Seed genes for ciliopathies were obtained from the Open Targets genetics portal, selecting those with gene-trait evidence scores > 0.5 for Genomics England PanelApp, Orphanet, Gene2Phenotype, UniProt literature, UniProt variants, ClinGen, and Gene Burden, and > 0.8 for ClinVar. Genes were curated to avoid overlap between subtypes of a disorder and to include recently identified genes. This resulted in the selection of 21 primary ciliopathies with at least two seed genes each. For mouse phenotypes, seed genes were also obtained from the Open Targets genetics portal, requiring a minimum of 10 seed genes per phenotype to exclude mouse models specific to only one monogenic disease. In total, 3,524 mouse phenotypes were included in the analysis.

We performed network propagation for ciliopathies and mouse phenotypes using the Personalized PageRank algorithm from the R package igraph v1.3.2, utilizing the same PPI network. Seed genes were assigned a weight of 1.

### Comparison of ciliopathies

We calculated the distance between ciliopathies through two methods. As a first method, the seed genes were used as binary input for computing the Jaccard distance between diseases using the R package stats v4.2.1. As a second method, z-scored network propagation scores for all 19,306 proteins were used as input for computing Euclidean distances between diseases. Using these distances, we hierarchically clustered the ciliopathies with Ward D2 linkage and compared the distances between ciliopathies with shared phenotypes by calculating the ROC curves and the AUROC. Ciliopathies were manually annotated with the following phenotypes: retinal defects, hearing loss, CNS defects, polydactyly, renal defects, and skeletal defects. We calculated the ROC curves and AUCs by considering ciliopathy pairs with at least one shared clinical phenotype as positives, as well as by taking each clinical phenotype separately.

### PPI network clustering

We clustered the PPI network using the Walktrap clustering algorithm from the R package igraph v1.3.2. Given that ciliary complexes are expected to be small, we performed re-clustering until every cluster consisted of <= 20 proteins or until 5 rounds of re-clustering were completed. We observed that protein modules with more than 20 proteins after 5 re-clustering rounds did not continue to split with further clustering attempts. To associate protein modules with ciliopathies and mouse phenotypes, we evaluated whether network propagation scores were significantly higher in the respective protein module compared to the full network using Wilcoxon Rank Sum tests. After p-value adjustment across all protein modules for each ciliopathy, we selected those with a p-value < 0.05 and at least one trait-related seed gene as associated with the trait. For naming of the protein modules, we conducted GO enrichment analysis combined with manual annotation. For visualization, we used Cystoscape [54] and annotated nodes with genes from SCGSv2 or CiliaCarta as ciliary and color-coded the genes based on their presence in ciliary protein complexes identified by Boldt et al. [33].

### Mouse phenotype clustering with ciliopathies

To compare mouse phenotypes with human ciliopathies, we calculated the Euclidean distance between the ciliopathies and all mouse phenotypes using scaled and centered network propagation scores. For each human ciliopathy, the 20 mouse phenotypes with the shortest distance were selected. Subsequently, we performed hierarchical clustering of the human ciliopathies with these selected mouse phenotypes using Euclidean distance and Ward D2 linkage. The number of clusters was determined to be 11 based on the optimal separation of human ciliopathies within the clusters. Mouse phenotypes were then annotated with their ancestor term and associated protein modules. For visualization, the ancestor terms and protein modules obtained from the network clustering with the highest fraction within each cluster were selected.

### Benchmarking ciliopathy gene prediction with mouse phenotype ranking

To test whether mouse phenotypes can contribute to candidate gene prediction for ciliopathies, we selected varying numbers of mouse phenotypes with the shortest Euclidean distances to each ciliopathy. Subsequently, we removed the known disease genes of the respective ciliopathies from the seed genes of the selected mouse phenotypes and reran the network propagation. The gene ranks based on network propagation scores were then combined from the selected mouse phenotypes using the geometric mean. Finally, we calculated the ROC curves and AUROCs for each ciliopathy with the known disease-causing genes as positives. It was found that 10 mouse phenotypes were optimal for recovering ciliopathy genes (Supplementary Fig. 1c). We also performed the same pipeline while selecting random mouse phenotypes and performed a t-test to compare the difference in AUCs.

### Gene expression of ciliopathy genes

To analyze the gene expression of ciliopathy genes, we obtained single-cell RNA sequencing data from HPA and centered and scaled the data across cell types. The data was split into a test and training set, with the test set containing 10% of known ciliopathy genes and 10% of all other genes, and the training set containing the remaining 90% of all genes. First, we performed an ANOVA with the training data to identify cell types with significantly different expression levels of known ciliopathy genes compared to the other genes. Cell types with a p-value < 0.01 were selected as features for a logistic regression model. The model’s weights were determined using the training data, with known ciliopathy genes as positives and the other genes as negatives. Finally, we evaluated the model by calculating the ROC curve and AUROC using the test data, treating the known ciliopathy genes as positives.

### Training and testing of model for candidate gene prioritization

To develop a model for prioritizing genes associated with different ciliopathies, we trained a logistic regression model incorporating three features: mouse phenotype ranking, network propagation p-values, and gene expression data. We selected four ciliopathies, each with at least 20 seed genes, to obtain model weights. For each ciliopathy, we performed multiple rounds of calculating separate feature scores based on five training genes, while the remaining disease genes were reserved as positives for the overall model. The number of rounds was determined by the total number of seed genes, ensuring that each gene was used only once in the training rounds. As negative controls, 150 genes not associated with the ciliopathy were randomly selected per round.

Network propagation scores were computed using the Personalized PageRank algorithm with the training genes as seed nodes. To prevent the high ranking of well-studied hub genes, we ran 1,000 permutations with random seed genes to compute p-values. For ranking of genes by mouse phenotypes, we calculated the Euclidean distances between scaled propagation scores derived from the five training genes and the scaled mouse phenotype propagation scores. The 10 mouse phenotypes with the shortest distance to the ciliopathy were selected, and ranks for every gene were combined using the geometric mean. For the gene expression scores, a logistic regression model, as described above, was trained in every round, excluding the test genes. The expression scores for each gene were then calculated using the trained model. All feature scores were min-max scaled to ensure they were in the same range.

After calculating the separate feature scores for each round and ciliopathy, the scores were aggregated to train one logistic regression model for all ciliopathies. This model used the test genes as positives and the randomly selected control genes as negatives, resulting in 727 positive genes and 3,643 negative genes. Separate models were trained with different combinations of the three features.

To evaluate our models for other ciliopathies, we selected six different ciliopathies with at least ten seed genes each. As with the training of the model, we used five seed genes as training genes and the remaining genes as test genes. In addition, 1,000 genes were randomly sampled as controls. Separate scores were again calculated using the training genes and combined using the trained models. The overall score was then used to calculate the PR AUC, partial AUROC with specificity between 0.95 and 1.0, and the AUROC. These metrics were compared between the different models to determine the optimal combination of features. This model was then used to calculate the final gene ranking for each ciliopathy, using all seed genes to calculate the separate feature scores.

### Assessment of gene prioritization

To gain insight into our gene prioritization, we conducted GSEA using various gene sets on the gene ranking excluding the known disease genes. Genes with gene-trait evidence scores below the thresholds defined for the seed genes were categorized into the ‘weak evidence’ gene set. In addition, genes from SCGSv2 and CiliaCarta were defined as ciliary genes. We also used localization data from HPA as gene sets. We performed GSEA for each ciliopathy separately and then counted the number of ciliopathies for which the gene sets (max size = 2000) were significantly (Benjamini-Hochberg adjusted p-value < 0.05) enriched among their high-ranking genes.

Furthermore, we examined the presence of high-ranking genes within protein modules. For each ciliopathy, we selected the top 100 ranked genes, excluding all seed genes. We calculated the fraction of these candidate genes in each protein module. As comparison, we also calculated the fraction of ciliary and known disease genes in each module, aggregated across all ciliopathies.

### Variant identification in patient cohort

A cohort of 75 patients with BBS (38), JBTS (13), and multi-systemic ciliopathies (24) was collected from the rare disease cohort in the Genomics England database. We scanned this cohort for homozygous and compound heterozygous variants in top candidate genes obtained from the gene prioritization and scanned these for their allele frequency and CADD scores. Among the findings, compound heterozygous variants in *CEP43* identified in an individual with a diagnosis of BBS, were found to be of particular interest. Following this discovery, we searched the broader Genomics England Database for variants in *CEP43*, leading to the identification of one more individual with an eye-related disorder carrying two mutations in *CEP43*. Lastly, a third individual was identified in a Saudi Arabian cohort through targeted reanalysis of existing sequencing data. For all coding variants, CADD, ESM1b, AlphaMissense, FoldX, and spliceAI scores were obtained to assess their potential impact. In addition, the AlphaFold prediction of CEP43 was retrieved from the database. Direct interactors of CEP43 and their association with human disorders and mouse phenotypes were retrieved and visualized to provide further contact to the identification of CEP43 as candidate gene.

## Data access

All data generated or analyzed during this study are included in this published article (and its Supplementary files). Publicly available repositories can be accessed as follows: OTAR interactome (https://ftp.ebi.ac.uk/pub/databases/IntAct/various/ot_graphdb/2022-07-22/), Open Targets Genetics portal (https://genetics.opentargets.org/), Experimental Factor Ontology (https://www.ebi.ac.uk/efo/), CiliaCarta (https://tbb.bio.uu.nl/john/syscilia/ciliacarta/), SCGSv2 (http://syscilia.org/), and HPA (https://www.proteinatlas.org/).

Research on the de-identified patient data used in this publication can be carried out in the Genomics England Research Environment subject to a collaborative agreement that adheres to patient led governance. All interested readers will be able to access the data in the same manner that the authors accessed the data. For more information about accessing the data, interested readers may contact or access the relevant information on the Genomics England website: https://www.genomicsengland.co.uk/research.

## Code availability

The network propagation and all subsequent analysis were performed using R software (v.4.2.1) as described in the methods. The following packages were used: igraph (v.1.3.2, for Personalized PageRank and walktrap clustering), pROC (v.1.18.0, for ROC curves and AUCs calculations when applicable), clusterprofiler (v.4.4.4, for GOBP enrichment analysis in the description of the modules as well as GSEA tests), ComplexHeatmap (v.2.12.1, for heatmap visualization), ggplot2 (v.3.4.0 for plotting), circlize (v.0.4.15) and RColorBrewer (v.1.1.3) both for color palette generation. sparklyr (v.1.7.8) and sparklyr.nested (v.0.0.3) both to obtain datasets, dplyr (v.1.1.4) and tidyverse (v.1.3.2) both to organize code, doParallel (v.1.0.17) and foreach (v.1.5.2) both for parallelization, dendextend (v.1.15.2 for dendogram plotting), ggpubr (v.0.5.0 for calculating statistics in plots), and org.Hs.eg.db (v.3.15.0 for gene ID mapping). Code is available on https://github.com/eaarts/networkPropagation/tree/main.

## Acknowledgements

E.M.A. is supported by Personalized Health and Related Technologies (PHRT) 2022-374. R.B.-G. received a grant from the Swiss National Science Foundation 310030_220012. P.B. is supported by the Helmut Horten Stiftung and the ETH Zurich Foundation. This research was made possible through access to data in the National Genomic Research Library, which is managed by Genomics England Limited (a wholly owned company of the Department of Health and Social Care). The National Genomic Research Library holds data provided by patients and collected by the NHS as part of their care and data collected as part of their participation in research. The National Genomic Research Library is funded by the National Institute for Health Research and NHS England. The Wellcome Trust, Cancer Research UK and the Medical Research Council have also funded research infrastructure.

## Author contributions

E.M.A., R.B.-G. and P.B. conceived and designed the project. E.M.A. performed most of the computational analysis with assistance from D.S.L.T. J.A.S., R.N., C.G.M., B.R., and R.E.A. performed the analysis in the Genomics England patient cohort and A.K., A.G., and M.H.A-H. provided patient information. E.M.A., R.B.- G., and P.B. wrote the manuscript with input from all authors.

## Supplementary Tables

**Supplementary Table 1:**
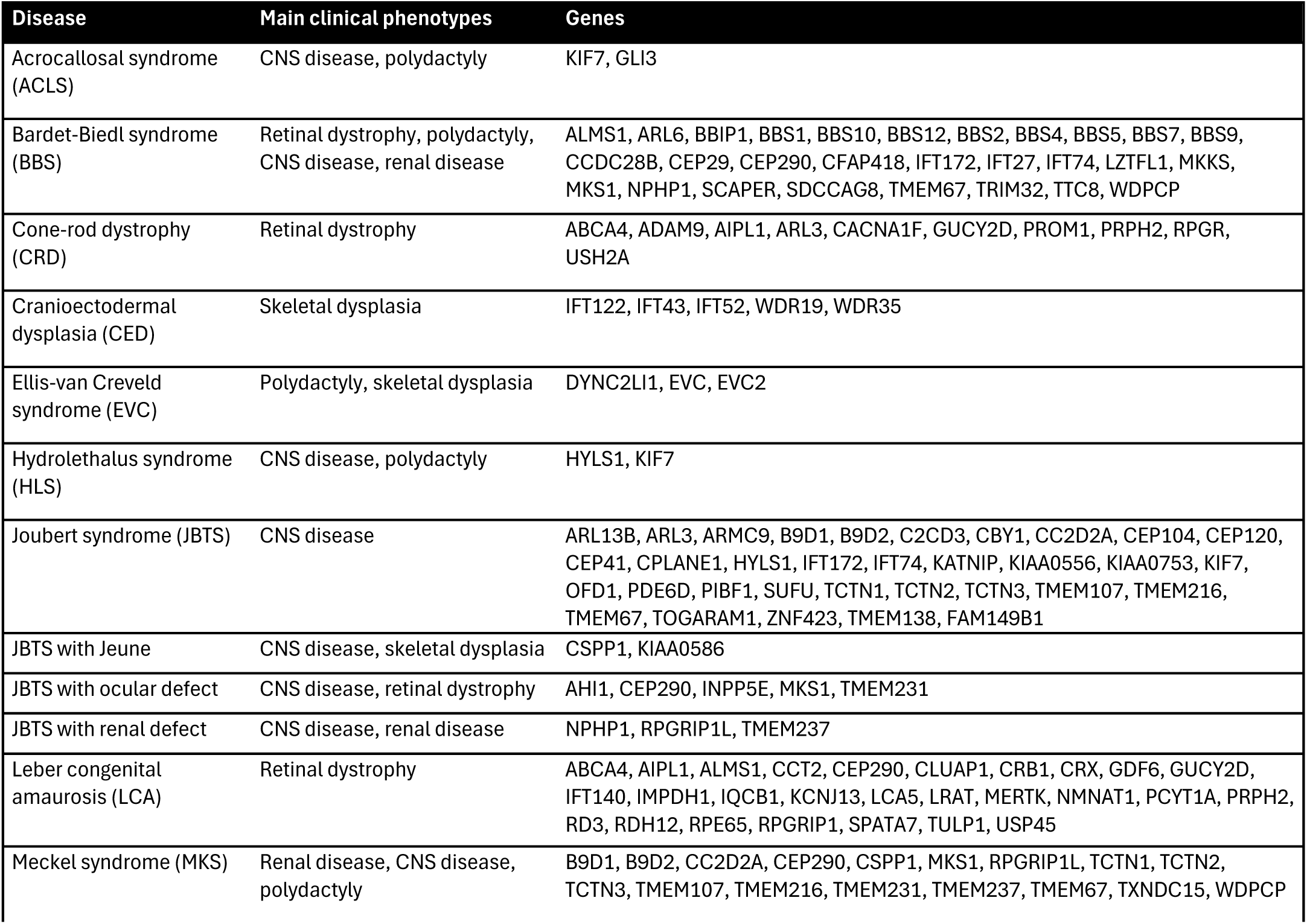
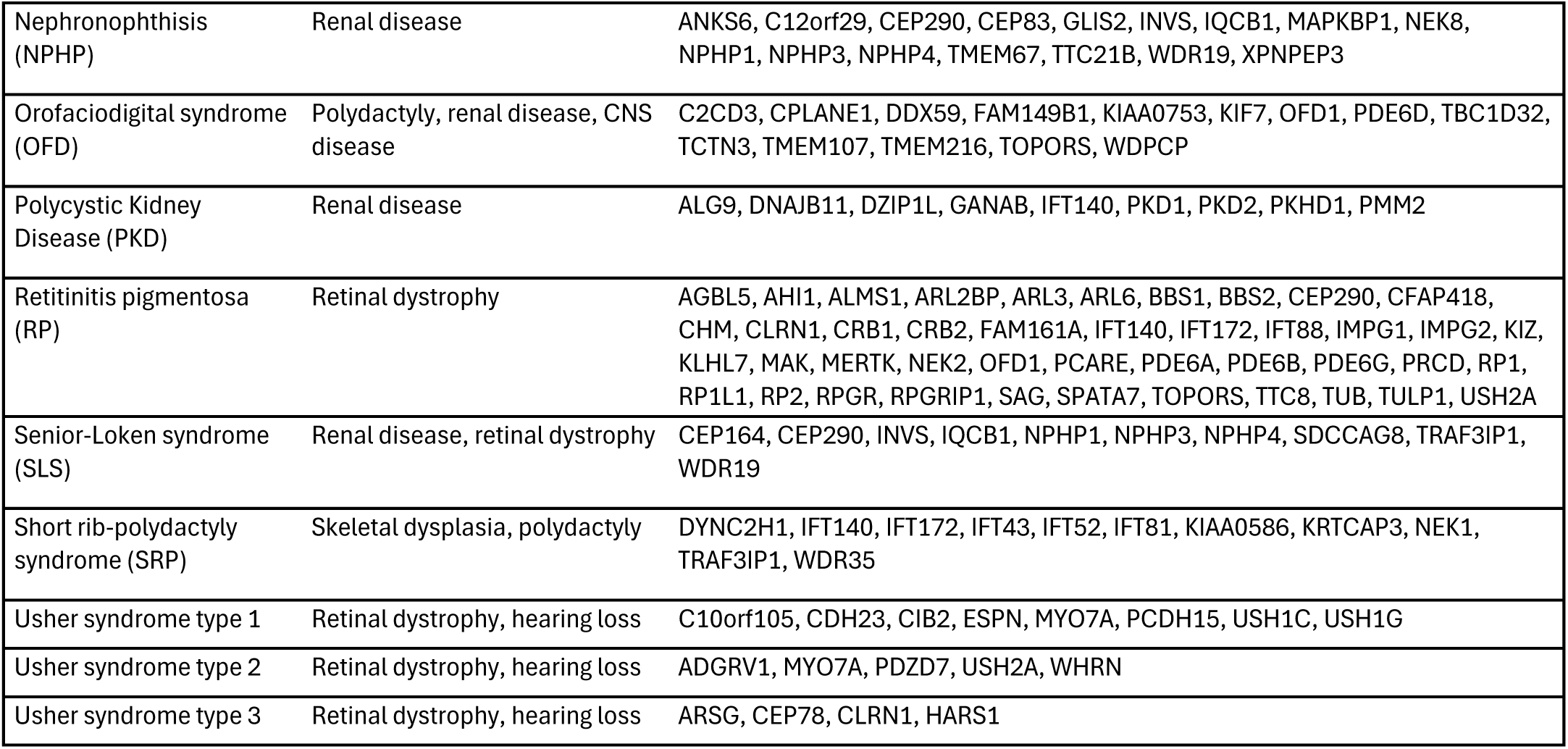
Clinical phenotypes and seed genes selected for each ciliopathy.

**Supplementary Table 10:** Identified CEP43 variants in individuals diagnosed with primary ciliopathies.

| Case ID | Age | Ethnicity | Consanguinity | CEP43 variants | Clinical diagnosis | Clinical phenotypes | Other genetic variants |
| --- | --- | --- | --- | --- | --- | --- | --- |
| 1 | 40-45 yrs | Caucasian | Yes | c.919+2T>G, p.(?) hmz | BBS | Mild progressive cone-rod dystrophy, postaxial polydactyly, congenital dysplasia of hip; Mild ID, obesity | N/A |
| 2 | 45-50 yrs | Asian | No | c.215G>A, p.(Arg72His), c.808A>G, p.(Arg270Gly) | Cone-rod dystrophy | Cone-rod dystrophy | CNGA3, c.952G>A p.Ala318Thr hmz |
| 3 | fetus | Saudi Arabian | Yes | c.35A>T, p.(Glu12Val) hmz | Skeletal dysplasia | Skeletal dysplasia | N/A |

## Supplementary Figures

**Supplementary Figure 1:**
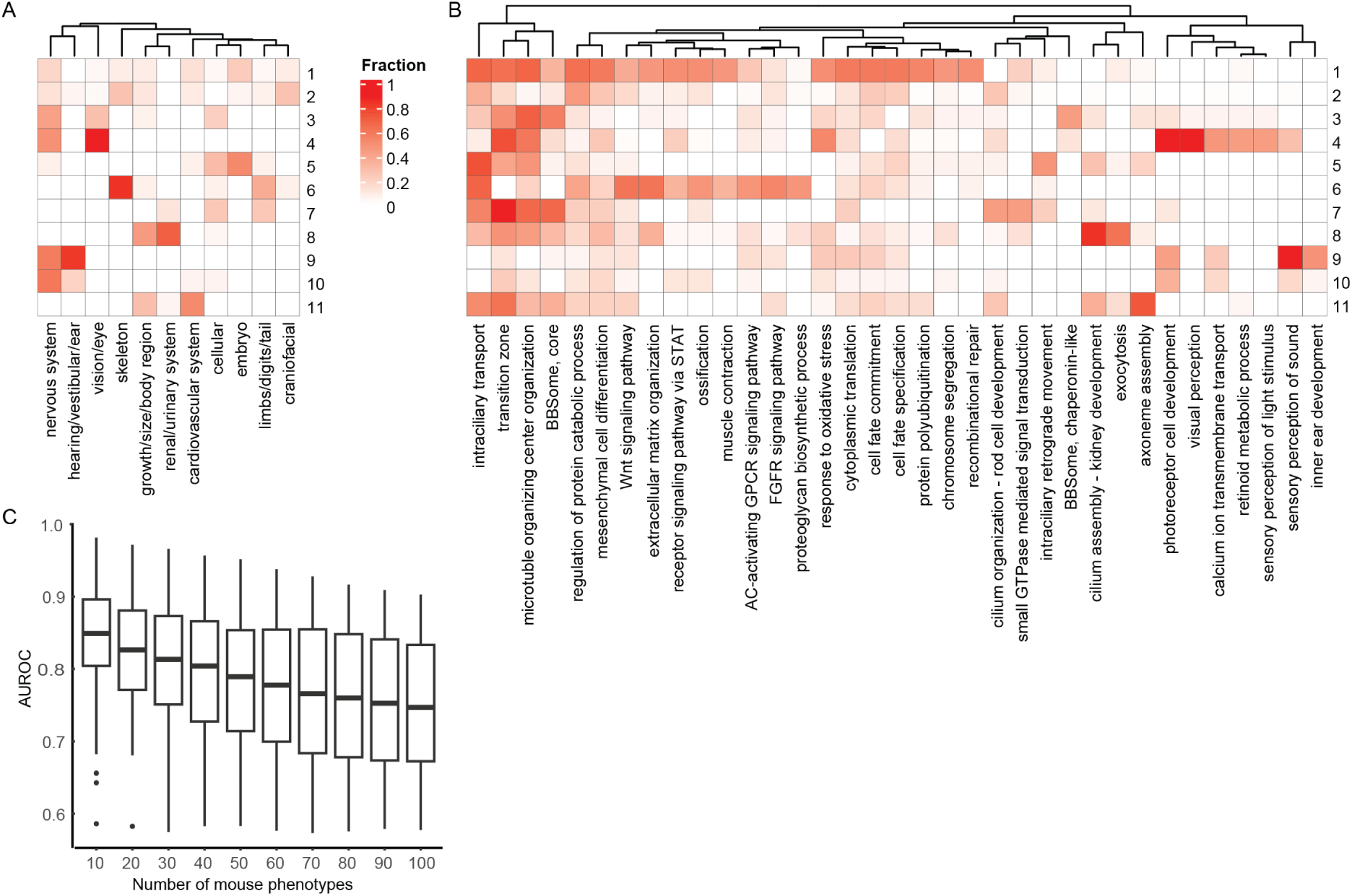
Mouse phenotype and human ciliopathy clusters associate to specific global phenotypes and protein modules. (A) Fraction of mouse phenotypes within each cluster from fig. 3a belonging to respective global phenotype, i.e., ancestor. (B) Fraction of mouse phenotypes within each cluster from fig. 3a associated with the respective protein module. (C) AUROC scores (median +/- IQR) for prediction of human ciliopathy genes with different numbers of mouse phenotypes with most similar network propagation scores to respective human ciliopathy. Ten mouse phenotypes give the highest AUROC scores for retrieving known human ciliopathy genes.

**Supplementary Figure 2:**
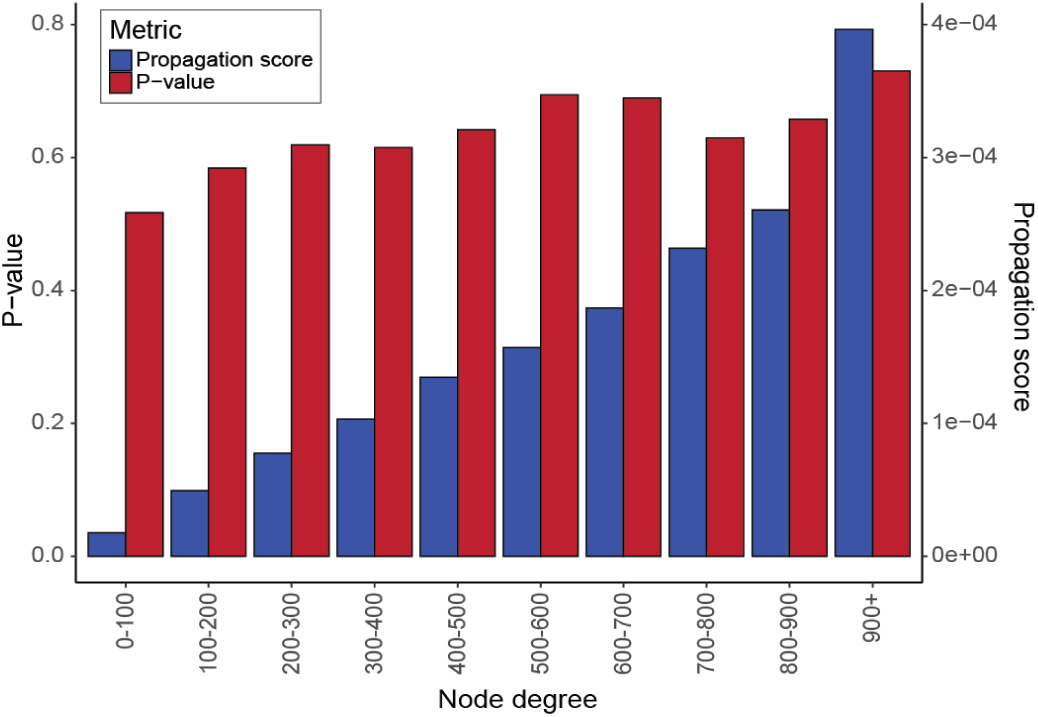
Network propagation scores depend on node degree. Per category of node degrees, the mean propagation scores (blue) and -log10 p-values (red) are visualized. P-values are obtained from permutation tests with randomly selected seed genes. Permutation removes correlation between node degree and network propagation scores.

**Supplementary Figure 3:**
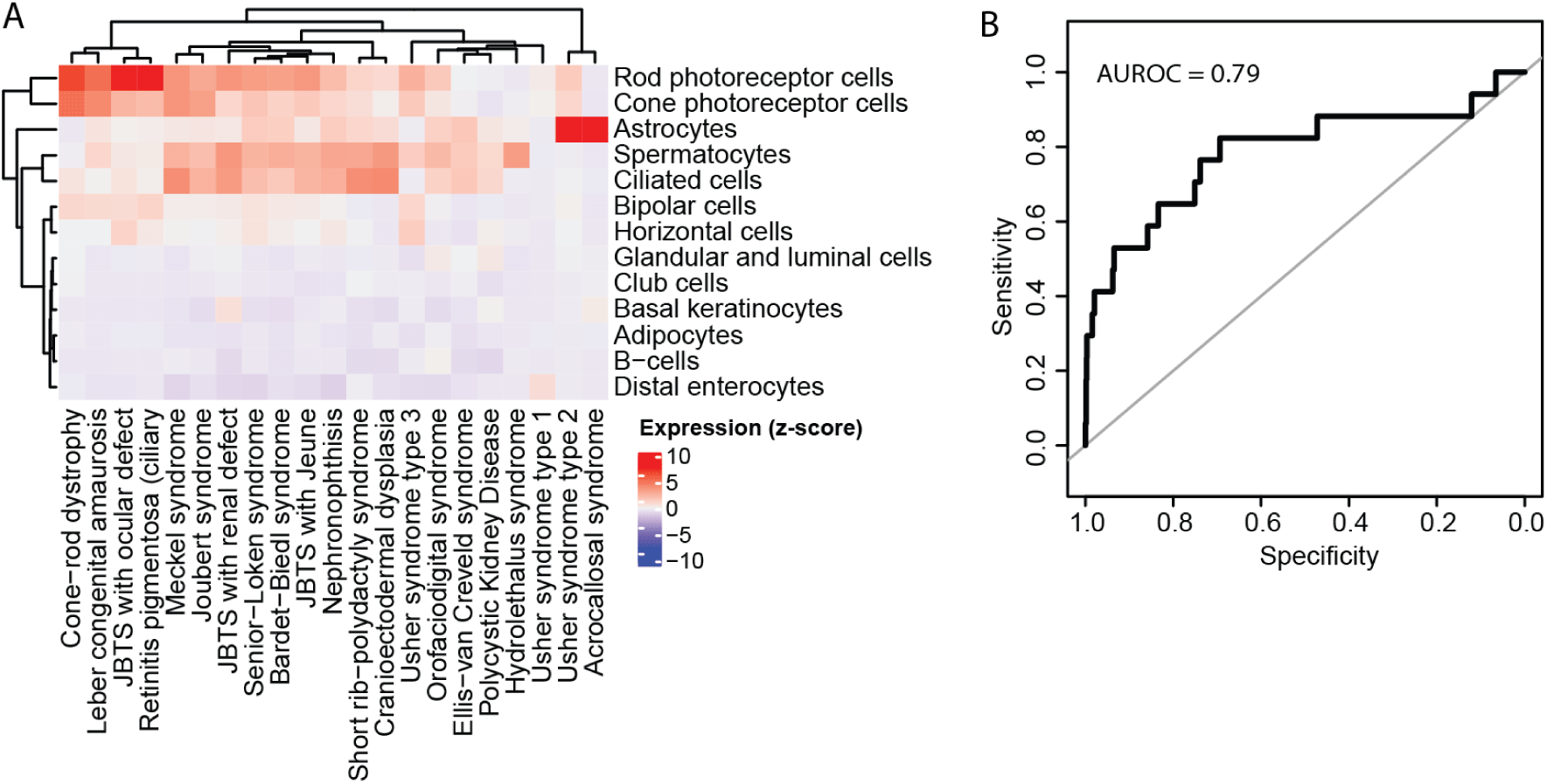
Gene expression to predict human ciliopathy genes. (A) Mean expression of ciliopathy-associated genes in selected cell types (z-scored over all cell types) obtained from HPA. Cell types were selected based on significant differential expression between known ciliopathy genes and other genes. (B) ROC curve for retrieval of human ciliopathy genes using a logistic regression model with gene expression in cell types from (A) as features.

**Supplementary Figure 4:**
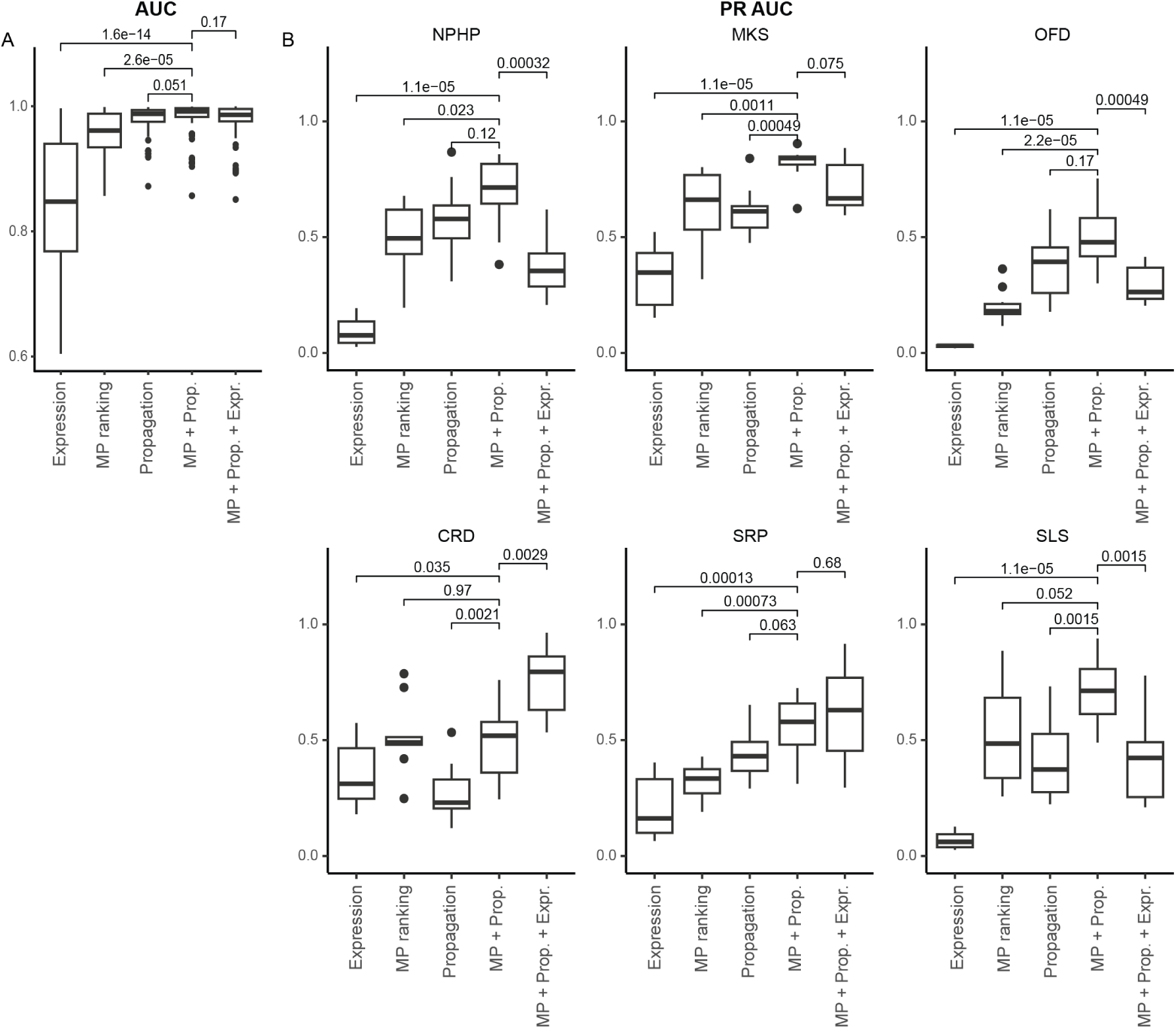
Accuracies of ciliopathy candidate gene prediction models. (A) AUC of models combining mouse phenotype ranking (MP), permuted network propagation scores (Propagation), and gene expression scores (Expression) in various combinations. (B) PR AUC of models in (A), split by ciliopathies used for testing the models. Barplots represent medium +/- IQR and t-tests for statistics.

**Supplementary Figure 5:**
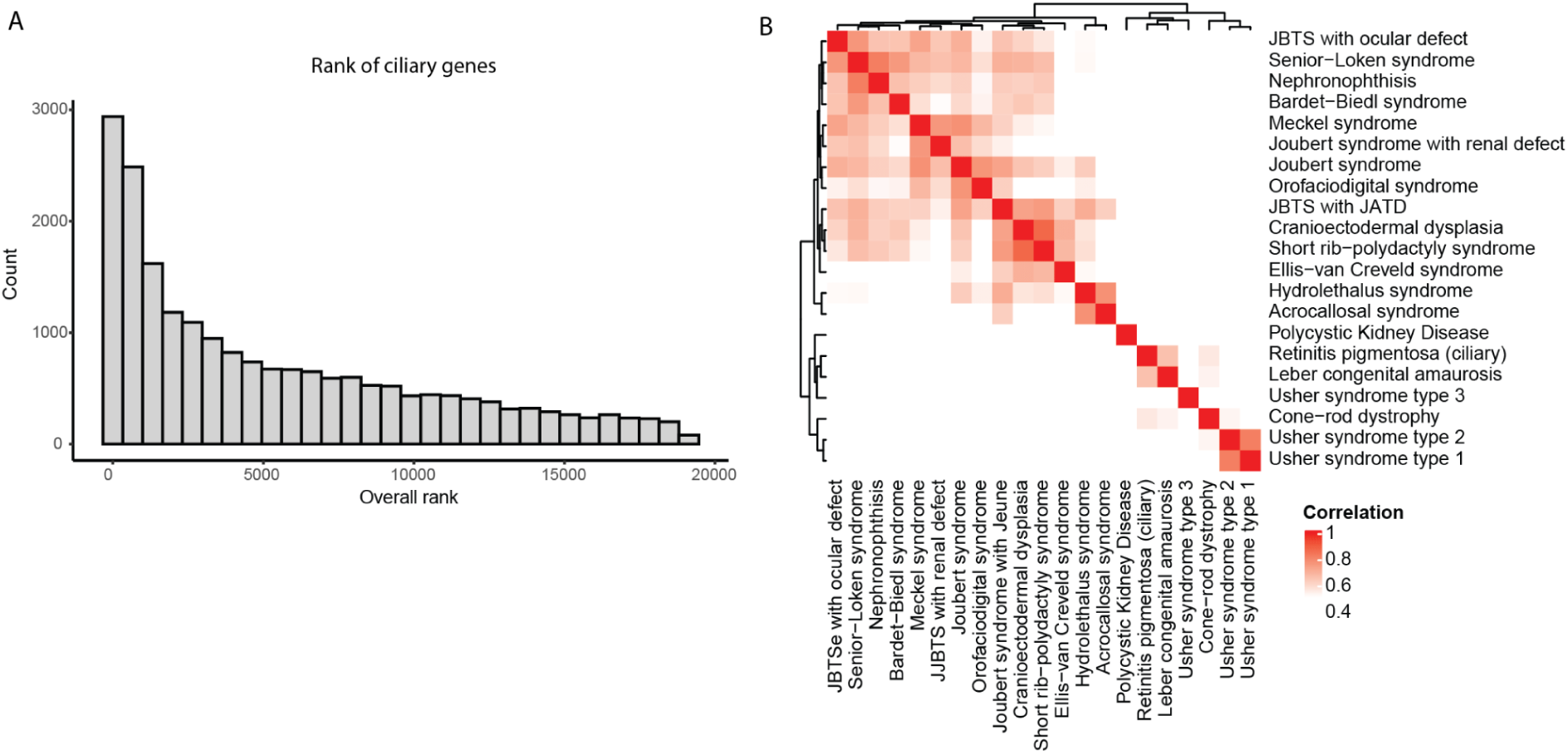
Scoring of genes per ciliopathy. (A) Overall rank from candidate gene prediction model of genes in CiliaCarta and/or SCGSv2 (ciliary genes) for 21 ciliopathies. Ciliary genes are more often ranked high but many are also ranked low. (B) Correlations of prediction scores between the 21 ciliopathies. The color scale starts at 0.5 to enhance visualization of correlation scores.

